# A simple method to genetically differentiate invasive F_1_ *Typha* hybrids (*T*. × *glauca*) and advanced-generation/backcrossed hybrids from parent species (*T. latifolia* and *T. angustifolia*) in eastern Canada and northeastern USA

**DOI:** 10.1101/2023.09.26.559561

**Authors:** Avery Chambers, Braidy Chambers, Deven Bhargava, Alberto Aleman, Marcel Dorken, Joanna Freeland

## Abstract

A widespread *Typha* (cattail) hybrid zone in southeastern Canada and northeastern USA comprises *T. latifolia, T. angustifolia*, and their hybrid *T*. × *glauca*. In this region the hybrid is considered invasive because of its detrimental impacts on wetlands and biodiversity. Researchers and managers are increasingly interested in tracking and controlling the spread of the invasive hybrid, but this can be hampered by challenges in differentiating taxa: while the parent species and F1 hybrids can often be identified from morphological characteristics, advanced-generation and backcrossed hybrids can be common, and these have obscured morphological distinctions among taxa. Here we present five species-specific genetic markers that differentiate *T. latifolia, T. angustifolia*, F_1_ hybrids, and advanced-generation/backcrossed hybrids. Unlike the pre-existing species-specific microsatellite markers, these markers require only PCR or PCR followed by a restriction enzyme digest, and can be visualized on an agarose gel. As a result, they require less specialized equipment than existing species-specific markers, and should therefore be useful for a larger group of practitioners. Future use of these markers could include monitoring range expansions, assessing the outcomes of restoration programs, and avoiding the unnecessary elimination of native *Typha*.

## Introduction

In regions around the Laurentian Great Lakes (LGL) and St. Lawrence Seaway a widespread *Typha* (cattail) hybrid zone comprises native *T. latifolia* (broad-leafed cattail), *T. angustifolia* (narrow-leafed cattail), and their hybrid *T. × glauca* (Smith 1967b; Grace and Harrison 1986b; Angeloni et al. 2006; Travis et al. 2010a; Kirk et al. 2011c; Ciotir and Freeland 2016; Pieper et al. 2020a).Here, *Typha × glauca* typically outcompetes both parental species to dominate wetlands (Travis et al. 2010b; Freeland et al. 2013a; Pieper et al. 2020a), and is considered invasive because it reduces biodiversity and alters wetland functions (Angeloni et al., 2006; Bansal et al., 2019; Boers et al., 2007; Farrer and Goldberg, 2014; Larkin et al., 2012; Lawrence et al., 2017a, 2016a; Tuchman et al., 2009a). As a result, conservation and land managers are increasingly tracking the composition of *Typha* stands to better understand invasion dynamics (Beas et al., 2013; Lawrence et al., 2017b; Lishawa et al., 2014; Mitchell et al., 2011; Tuchman et al., 2009b), and managing *Typha* stands to reduce dominant taxa including *T*. x *glauca* (Bansal et al., 2019; Lawrence et al., 2016b; Lishawa et al., 2017; Solberg and Higgins, 1993). However, inconclusive identification of *T*. × *glauca* can hamper these goals.

In the North American hybrid zone, asymmetric hybridization involving maternal *T. angustifolia* and paternal *T. latifolia* produces F_1_ *T*. × *glauca* (Ball & Freeland, 2013; Pieper et al., 2017). F_1_ hybrids can backcross bidirectionally with *T. angustifolia* and asymmetrically with *T. latifolia* (the latter being the pollen donor), and can also create viable advanced-generation hybrids; backcrossed and advanced-generation hybrids (BC/AG hybrids) are fertile, although F_2_ hybrids have shown evidence of hybrid breakdown (Bhargav et al., 2022; Pieper et al., 2017). Hybrid *Typha* swarms therefore comprise parent species, F_1_ hybrids, and advanced-generation/backcrossed hybrids (Freeland et al., 2013b; Kirk et al., 2011; Pieper et al., 2020b; Travis et al., 2010a). BC/AG hybrids complicate taxon identification on the basis of morphological characters. *Typha latifolia* and *T. angustifolia* can often be differentiated on the basis of leaf width, presence or absence of a gap between staminate and pistillate spikes, and spike width (Bansal et al., 2019; Finkelstein, 2003; J. Grace & Harrison, 1986; Kirk et al., 2011b; Krattinger, 1975; S. Smith, 1967; Snow et al., 2010). However, while F_1_ hybrids have roughly intermediate leaf widths and flower gaps (J. Grace and Harrison, 1986; Kirk et al., 2011; Smith SG, 2020; Snow et al., 2010), BC/AG hybrids lead to considerable overlap between these measurements in hybrids versus parental species (Geddes et al., 2021b; Tangen et al., 2022b).Furthermore, nitrogen levels impact leaf width in *T. latifolia, T. angustifolia*, and *T*. x *glauca*, and leaf width in *T*. × *glauca* further differs between flowering versus non-flowering shoots (Shapro, 2022). In another study shoot dimensions and microscopic analysis of leaves were used to differentiate the three taxa, but this was partly based on 14 hybrid plants that could have been F_1_s or AG/BC hybrids (Wasko et al. 2022), and therefore the widespread applicability of this method requires additional validation.

Flowering traits have also been used to provisionally identify *T. latifolia, T. angustifolia*, and *T*. × *glauca*, although the flowering season is relatively short, and in any given year many ramets – and even entire stands -do not flower. While pollen grains are tetrad in *T. latifolia*, monad in *T. angustifolia*, and a mixture of monads, dyads, triads and tetrads in hybrids (Dugle and Copps, 1972; Finkelstein, 2003b), we have found that this method does not always allow for accurate differentiation between *T. angustifolia* and hybrids (based on subsequent genotyping methods; see below), and this method is further limited by the short pollen-shedding season (e.g. approximately two weeks across populations in southern Ontario). Bracteoles have been used to differentiate parental species from F_1_ hybrids: *T. latifolia* lacks bracteoles, *T. angustifolia* has dark bracteoles that are wider than adjacent stigmas, and *T*. × *glauca* has bracteoles that are lighter in colour and narrower than adjacent stigmas (Smith SG, 2020). However, there have been no investigations of bracteoles in AG/BC hybrids, nor have bracteoles been compared from mature flowers; additionally, as noted above, *Typha* stands do not always include flowering shoots.

Genotyping can provide a more exact method of taxon identification than morphology, particularly when there are cryptic AG/BC hybrids or no flowering plants. The most common method that has been used to genetically differentiate *T. angustifolia, T. latifolia*, and *T*. × *glauca* is microsatellite genotyping (e.g.(Bhargav et al., 2022; Freeland et al., 2013b; Pieper et al., 2020b, 2018; Tisshaw et al., 2020; Travis et al., 2010a; Zapfe and Freeland, 2015): four microsatellite loci generate species-specific alleles in southeastern Canada and northeastern USA (Kirk et al., 2011; Snow et al., 2010). While this method is reliable, microsatellite genotyping requires equipment and expertise that is often beyond the scope of many field biologists. We have developed a set of markers that genetically discriminate *T. angustifolia, T. latifolia*, and hybrids (F_1_s versus advanced-generation/backcrossed hybrids), and which require less specialized and hence more widely accessible laboratory equipment: with taxon-based genetic differences apparent following PCR and gel electrophoresis, or following PCR and a restriction enzyme digest (revealing a restriction fragment-length polyorphism, PCR-RFLP) and gel electrophoresis.

## Methods

### Marker development

PCR-RFLP markers are species-specific if the PCR primers anneal to regions that are conserved between species, and the amplified region includes a restriction enzyme recognition site that depends on a single nucleotide polymorphism (SNP) for which alternative alleles are fixed in each species; as a result, when the amplified fragment is incubated with a restriction enzyme (RE), it will be either cut into two fragments or left uncut depending on the species. Provided the fragments are of appropriate size (see below), cut versus uncut amplicons can be differentiated on an agarose gel. In hybrid zones that comprise solely parent species and F_1_ hybrids, a single species-specific PCR-RFLP marker (i.e. one locus) is sufficient to differentiate parents from hybrids, because it will digest the amplified DNA of one but not the other parent, and the F_1_ hybrids will be heterozygous for cut and uncut alleles; this approach was taken to differentiate two subspecies of *Phragmites australis* and their hybrid (Wendell et al., 2021). However, the situation is more complicated in the *T. angustifolia – T. latifolia* hybrid zone because at any given locus an AG/BC hybrids could have two alleles from one or the other parent. We therefore need data from multiple species-specific loci before concluding that an individual is a parent species versus an AG/BC hybrid: a minimum of four loci are needed in order to differentiate the parents and the main categories of hybrids (F1, AG/BC) in a hybrid zone (Boecklen and Howard, 1997).

In order to identify species-specific SNPs that could be incorporated into PCR-RFLP markers, we used genome-wide sequences of 12 *T. angustifolia* from Ontario, Quebec, and Nova Scotia Canada, and Michigan, USA, and 44 *T. latifolia* from British Columbia, Manitoba, Ontario, Quebec, New Brunswick, and Nova Scotia, Canada, and Maine, Michigan, and Florida, USA (Aleman et. al., 2023). For each sample, we used mapped and indexed nuclear data to produce a consensus sequence with degenerate nucleotides in SAMtools 1.15.1 (Li et al., 2009), requiring minimum mapping and sequencing qualities of 20, allowing for indels, setting the ploidy as 2 and missing data as N. We aligned all the sequences by applying MAFFT 7.0 default settings (i.e., using the flag *--nzero*) (Katoh et al., 2019). We identified species-specific SNPs (fixed for the same allele within species and alternate alleles between species) in 1.5 Kbp windows and then searched 20 bp sequences up- and downstream of each SNP for RE recognition sites using NEBcutter V2.0 (New England BioLabs; http://nc2.neb.com/NEBcutter2/index.php). When we determined that a species-specific SNP dictated the presence/absence of an RE recognition site, we scanned the sequence 1,500 bp upstream and 1,500 bp downstream with GenScript Restriction Enzyme Map Analysis Tools (https://www.genscript.com/tools/restriction-enzyme-map-analysis) to ensure that the sequence contained only one recognition site for that RE. When that criterion was met, we analyzed the 3,000 bp sequence with Primer Blast (Ye et al., 2012) to design primer pairs that would anneal to conserved regions in both species, amplify a fragment between ∼500 bp and 1500 bp, and generate post-RE digest fragment sizes of at least 100 bp. This left us with 20 loci for further testing.

### Marker validation

We tested the primers and enzymes on *Typha* DNA that had been assigned to taxon in previous studies on the basis of species-specific microsatellite alleles (Bhargav et al., 2022; Pieper et al., 2020b; Tisshaw et al., 2020). We included samples previously identified as *T. latifolia, T. angustifolia*, and F1 *T*. × *glauca* spanning a wide geographical region: in total we genotyped 16 *T. angustifolia* from Ontario and Nova Scotia, Canada, and Michigan, USA; 16 *T. latifolia* from Ontario, New Brunswick, and Nova Scotia, Canada, and Michigan, USA; and 21 F_1_ hybrids from Ontario and Nova Scotia, Canada, and Michigan, USA (Figure 1). We first used gradient PCRs to identify the optimal annealing temperature, and then tested the results of RE incubation for the loci that consistently produced PCR products of the anticipated size. Sixteen of the PCR-RFLP 20 markers developed *in silico* were excluded for various reasons (e.g. non-specific priming, RE digests that were not species-specific), and detailed methods are therefore provided only for the four loci that generated repeatable, species-specific patterns. For these loci, the amplification reactions included 12.5 uL 2 × Froggabio master mix, either 0.4 mM (AseI, BsrI) or 0.2 mM (BtsCI, PhyC) of forward and reverse primers, 1 uL DNA, and ddH_2_O for a final volume of 25 mL. Cycling conditions comprised an initial denaturation of 94 °C for two minutes; 35 cycles of 45 sec at 94 °C, 45 sec annealing (Table 1), and 60 sec at 68 °C; and a final extension of 68 °C for two minutes. PCR amplification was verified on a 1% agarose gel with a 100 bp ladder (Froggabio) for reference.

**Fig. 1.**
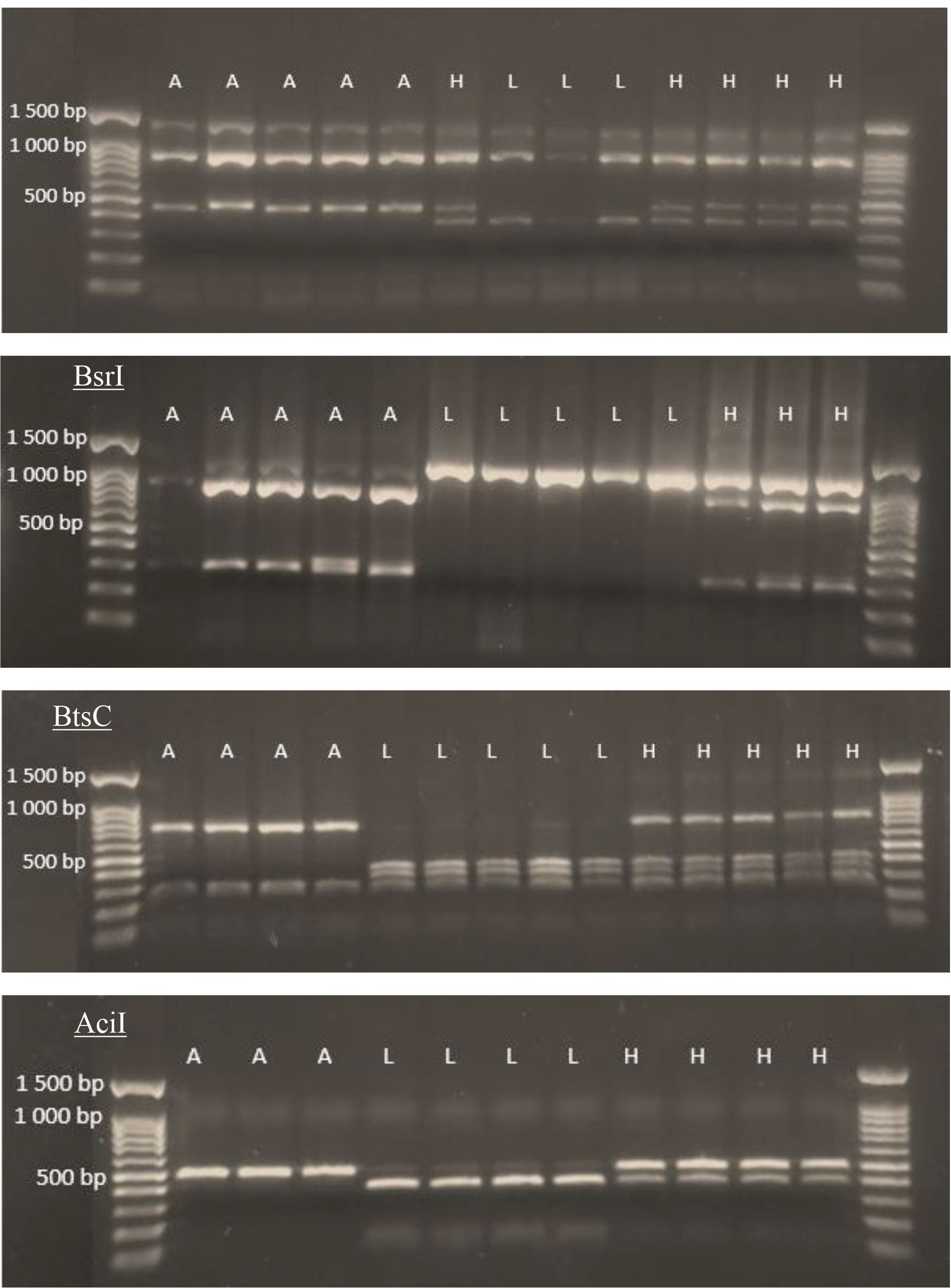
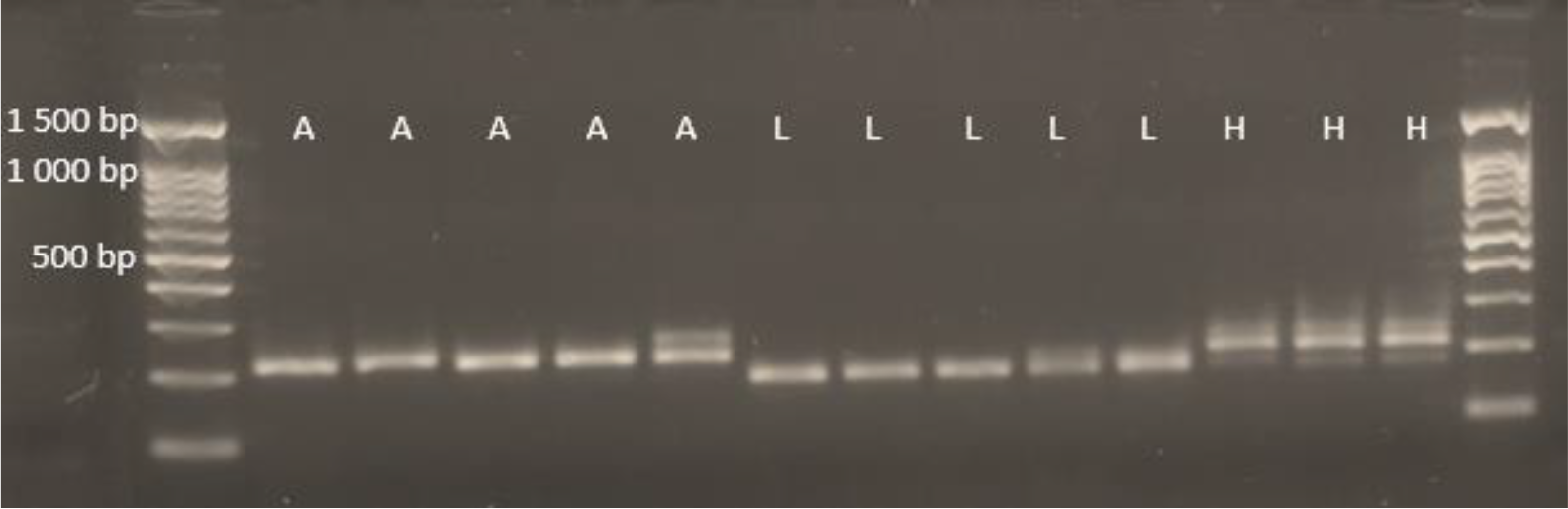
Gel images depicting the species-specific PCR-RFLP polymorphisms (AseI, BsrI, BstCI, Acil) or the PCR allele sizes (TA3). “A” = *Typha angustifolia*, “L” = *Typha latifolia*;,”H” = *Typha* × *glauca*. Fragment lengths are compared to 100 base pair (bp) DNA ladders.

**Table 1.**
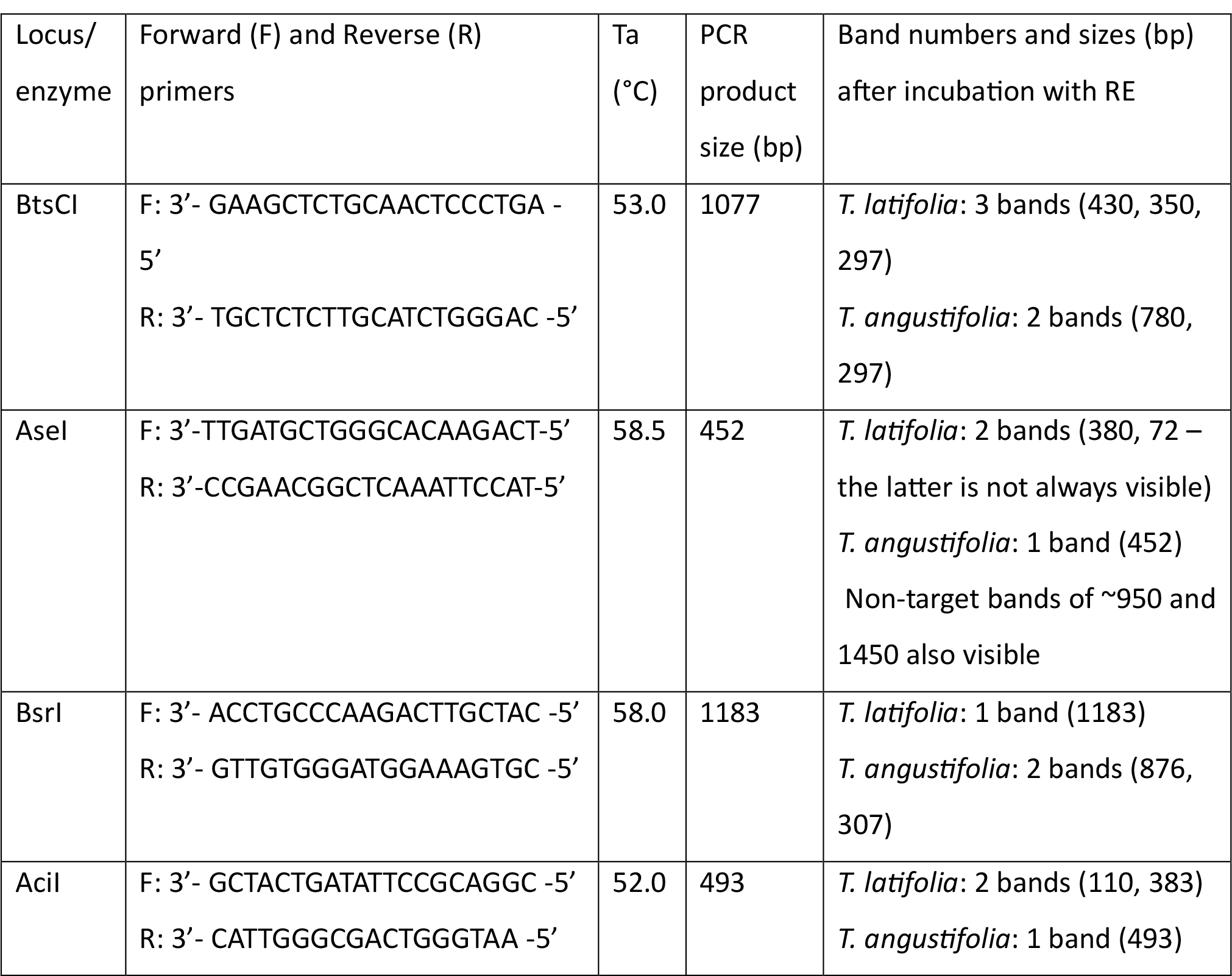
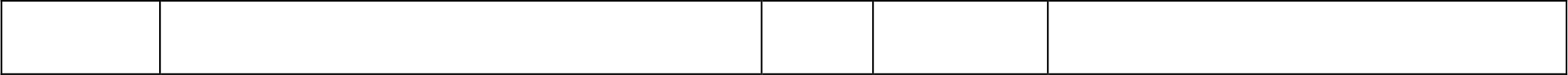
Locus name (named after the with which the PCR product was incubated), primer sequences, annealing temperature (Ta), size of amplified product, and expected band sizes following RE incubation.

Successful amplifications were incubated with the relevant RE. All enzymes were from New England Biolabs. Each RE incubation was done in a 10uL reaction with 1 × rCutSmart buffer, either 3 uL (AciI), 4 uL (NheI, BtsCI, AseI) or 8 uL (BsrI) of PCR product (varied according to the intensity of PCR bands), either 5U (AciI BsrI, AseI) or 8U (BtsCI) of enzyme, and ddH2O to a final volume of 10 uL. Incubation temperatures and inactivation times and temperatures followed the manufacturer’s protocols, although we increased the incubation time to 1.5 hours because the shorter recommended incubation time sometimes resulted in incomplete digests. After inactivation, samples were visualized on a on a 1% agarose gel with a 100 bp ladder (Froggabio) for reference. Table 1 shows the primers, PCR conditions, enzymes, and expected band sizes for the five PCR-RFLP markers.

In addition to the four PCR-RFLP markers described above, we tested the efficacy of an agarose gel-based screening of the microsatellite locus TA3. We chose this locus because previous studies have shown that in most cases *T. angustifolia* and *T. latifolia* alleles at TA3 differ by at least 30 bp, the largest size difference that has been found between species-specific microsatellite alleles (H Kirk et al., 2011; Snow et al., 2010). We therefore investigated whether this size difference was sufficient to differentiate species on an agarose gel, potentially removing the need for microsatellite electropherograms. We amplified microsatellite alleles at locus TA3 following the methods of (Kirk et al. 2011b). Because there could be as few as 30 bp between amplified alleles, we ran the PCR products on a 2% agarose gel to improve separation.

## Results and Discussion

Four of our PCR-RFLP markers generated species-specific patterns (Table 1, Figure 2) across all of the geographical regions that we sampled. The TA3 primer pair (the microsatellite locus) typically generated one band for each of *T. latifolia* and *T. angustifolia* that are consistent with the size ranges of 172-184 bp and 215-221 bp, respectively, that are reported in (Kirk et al., 2011). In one of the *T. angustifolia* samples there was a second, slightly larger band (Figure 1), which is not unexpected when amplifying regions of high repeat (comparable to the ‘stutters’ in microsatellite electropherograms). The hybrid samples have three TA3 bands corresponding with the *T. latifolia* and *T. angustifolia* allele sizes plus the ‘extra’ *T. angustifolia* band (which is clearly different from the *T. latifolia* band). In the hybrids, the *T. latifolia* band is consistently fainter than the *T. angustifolia* band, and while this marker can be considered diagnostic on a 2% agarose gel, the overall proximity of the bands mean that it will be important to include positive controls for TA3, i.e. amplifications from known *T. angustifolia, T. latifolia*, and *T*. × *glauca* DNA.

We have characterised a total of five species-specific markers that require PCR and – for four of the markers -incubation with a RE, followed by visualization of results on an agarose gel. This method is cheaper and faster than microsatellite genotyping, requires less equipment and expertise, and can be used on all plants that are sampled during the growing season.

Furthermore, we have demonstrated its reliability across a large geographical range including areas around the Laurentian Great Lakes, St. Lawrence Seaway, and maritime Canada. Potential applications include differentiation of F_1_ from AG/BC hybrids to better understand the successful propagation and spread of different hybrid classes; differentiation of *T. angustifolia* from *T*. × *glauca* to investigate whether the recently documented range expansion of the latter (Tangen et al. 2021) extends beyond that of the former; and identification of native *T. latifolia* from invasive hybrids for the purpose of effectively managing and restoring invaded wetlands. The method described here should therefore be useful to a relatively large group of researchers and managers.

## Acknowledgements

Funding for this project was provided by Discovery Grants from the Natural Sciences and Engineering Research Council of Canada awarded to M. Dorken (RGPIN-2018-04866) and J. Freeland (RGPIN-2017-04371). A. Chambers, B. Chambers, and D. Bhargav received NSERC USRA summer scholarships. Many thanks to Sara Pieper, Kat Tisshaw, and Vikram Bhargav for DNA samples of known taxon.

---

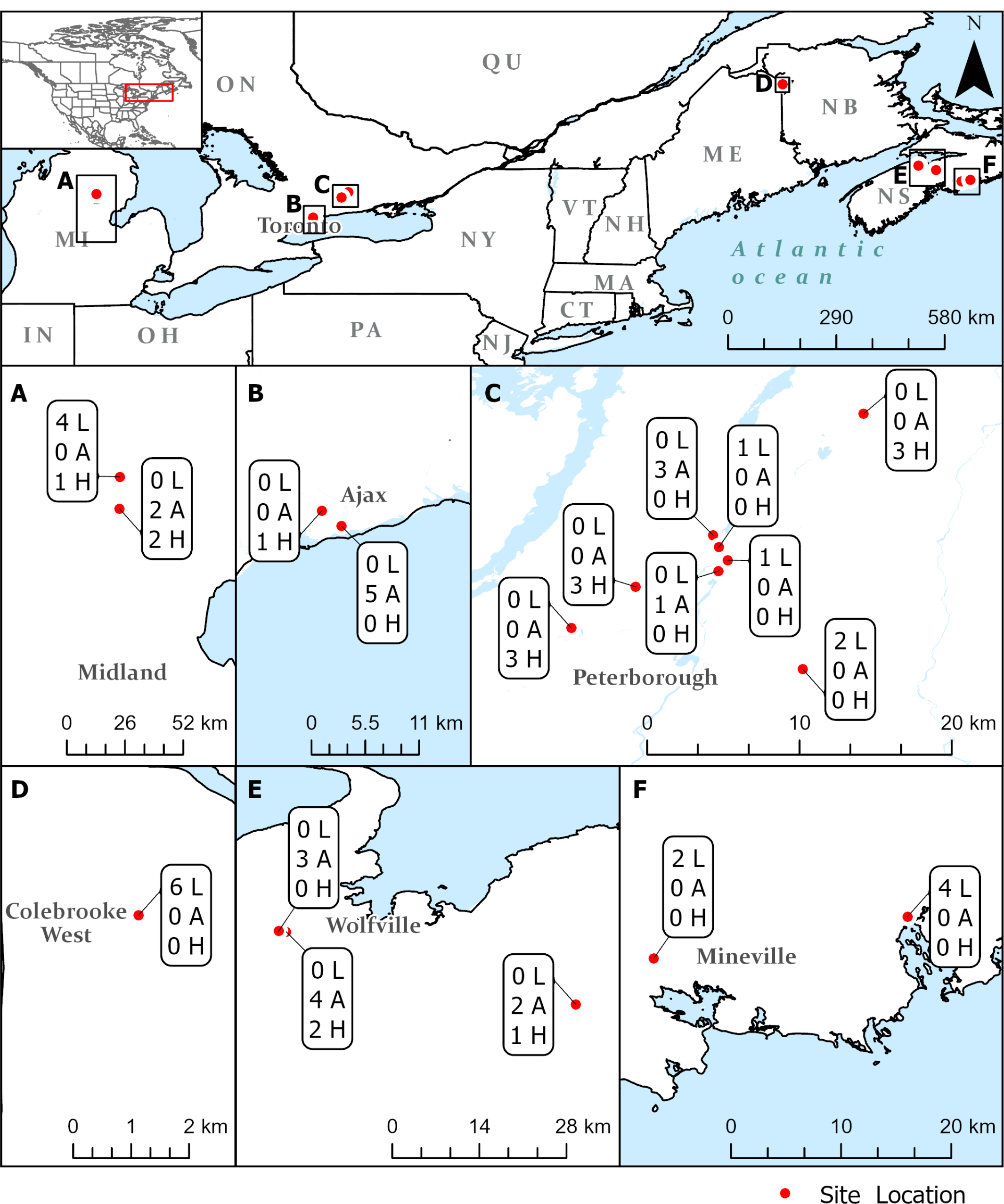

## Literature cited

Angeloni, N.L., Jankowski, K.J., Tuchman, N.C., Kelly, J.J., 2006. Effects of an invasive cattail species (Typha x glauca) on sediment nitrogen and microbial community composition in a freshwater wetland. FEMS Microbiol Lett 263. 10.1111/j.1574-6968.2006.00409.x

Ball, D., Freeland, J.R., 2013. Synchronous flowering times and asymmetrical hybridization in Typha latifolia and T. angustifolia in northeastern North America. Aquat Bot 104. 10.1016/j.aquabot.2012.08.006

Bansal, S., Lishawa, S.C., Newman, S., Tangen, B.A., Wilcox, D., Albert, D., Anteau, M.J., Chimney, M.J., Cressey, R.L., DeKeyser, E., Elgersma, K.J., Finkelstein, S.A., Freeland, J., Grosshans, R., Klug, P.E., Larkin, D.J., Lawrence, B.A., Linz, G., Marburger, J., Noe, G., Otto, C., Reo, N., Richards, J., Richardson, C., Rodgers, L.R., Schrank, A.J., Svedarsky, D., Travis, S., Tuchman, N., Windham-Myers, L., 2019. Typha (Cattail) Invasion in North American Wetlands: Biology, Regional Problems, Impacts, Ecosystem Services, and Management. Wetlands. 10.1007/s13157-019-01174-7

Beas, B.J., Smith, L.M., LaGrange, T.G., Stutheit, R., 2013. Effects of sediment removal on vegetation communities in Rainwater Basin playa wetlands. J Environ Manage 128. 10.1016/j.jenvman.2013.04.063

Bhargav, V., Freeland, J., Dorken, M., 2022. Evidence of hybrid breakdown among invasive hybrid cattails (Typha x glauca). Heredity (Edinb) 129, 195–201.

Boecklen, W.J., Howard, D.J., 1997. Genetic analysis of hybrid zones: Numbers of markers and power of resolution. Ecology 78. 10.1890/0012-9658(1997)078[2611:GAOHZN]2.0.CO;2

Boers, A.M., Veltman, R.L.D., Zedler, J.B., 2007. Typha×glauca dominance and extended hydroperiod constrain restoration of wetland diversity. Ecol Eng 29, 232–244. 10.1016/j.ecoleng.2006.04.011

Ciotir, C., Freeland, J., 2016. Cryptic intercontinental dispersal, commercial retailers, and the genetic diversity of native and non-native cattails (Typha spp.) in North America. Hydrobiologia 768, 137–150. 10.1007/s10750-015-2538-0

Dugle, J.R., Copps, T.P., 1972. Pollen characteristics of Manitoba cattails. Canadian Field Naturalist 86, 33–40.

Farrer, E.C., Goldberg, D.E., 2014. Mechanisms and reversibility of the effects of hybrid cattail on a Great Lakes marsh. Aquat Bot 116, 35–43.

Finkelstein, S.A., 2003a. Identifying pollen grains of Typha latifolia, Typha angustifolia, and Typha ×glauca 81, 985–990. 10.1139/B03-084

Finkelstein, S.A., 2003b. Identifying pollen grains of Typha latifolia, Typha angustifolia, and Typha × glauca. Canadian Journal of Botany 81, 985–990. 10.1139/b03-084

Freeland, J., Ciotir, C., Kirk, H., 2013a. Regional differences in the abundance of native, introduced, and hybrid Typha spp. in northeastern North America influence wetland invasions. Biol Invasions 15, 2651–2665. 10.1007/s10530-013-0481-4

Freeland, J., Ciotir, C., Kirk, H., 2013b. Regional differences in the abundance of native, introduced, and hybrid Typha spp. in northeastern North America influence wetland invasions. Biol Invasions 15, 2651–2665. 10.1007/s10530-013-0481-4

Geddes, P., Murphy, L., Astudillo-Scalia, Y., Blasini, D., Nugent, S., Ríos, M.J., Schirmer, A.E., Olfelt, J.P., 2021a. Microsatellite Markers Reveal Unprecedented High Frequencies of Hybridization among Typha Species in the Midwestern US. Wetlands 41, 24. 10.1007/s13157-021-01429-2

Geddes, P., Murphy, L., Astudillo-Scalia, Y., Blasini, D., Nugent, S., Ríos, M.J., Schirmer, A.E., Olfelt, J.P., 2021b. Microsatellite Markers Reveal Unprecedented High Frequencies of Hybridization among Typha Species in the Midwestern US. Wetlands 41. 10.1007/s13157-021-01429-2

Grace, J., Harrison, J., 1986. The biology of Canadian weeds: 73. Typha latifolia L., Typha angustifolia L. and Typha x glauca Godr. Canadian Journal of Plant Science 66, 361–379. 10.4141/cjps86-051

Grace, J.B., Harrison, J.S., 1986. The biology of Canadian weeds: 73. Typha latifolia L, Typha sngustifolia L and Typha X glauca Godr. Canadian Journal of Plant Science 66, 361–379. 10.4141/cjps86-051

Kirk, H., Connolly, C., Freeland, J.R., 2011. Molecular genetic data reveal hybridization between Typha angustifolia and Typha latifolia across a broad spatial scale in eastern North America. Aquat Bot 95, 189–193. 10.1016/j.aquabot.2011.05.007

Kirk, H., Connolly, C., Freeland, J.R., 2011. Molecular genetic data reveal hybridization between Typha angustifolia and Typha latifolia across a broad spatial scale in eastern North America. Aquat Bot 95. 10.1016/j.aquabot.2011.05.007

Kirk, H, Connolly, C., Freeland, J.R., 2011. Molecular genetic data reveal hybridization between Typha angustifolia and Typha latifolia across a broad spatial scale in eastern North America. Aquat Bot 95, 189–193. 10.1016/j.aquabot.2011.05.007

Krattinger, K., 1975. Genetic mobility in Typha. Aquat Bot 1, 57–70. 10.1016/0304-3770(75)90007-8

Larkin, D.J., Lishawa, S.C., Tuchman, N.C., 2012. Appropriation of nitrogen by the invasive cattail Typha×glauca. Aquat Bot 100, 62–66. 10.1016/j.aquabot.2012.03.001

Lawrence, B.A., Bourke, K., Lishawa, S.C., Tuchman, N.C., 2016a. Typha invasion associated with reduced aquatic macroinvertebrate abundance in northern Lake Huron coastal wetlands. J Great Lakes Res 42, 1412–1419. 10.1016/j.jglr.2016.08.009

Lawrence, B.A., Lishawa, S.C., Hurst, N., Castillo, B.T., Tuchman, N.C., 2017a. Wetland invasion by Typha×glauca increases soil methane emissions. Aquat Bot 137, 80–87. 10.1016/j.aquabot.2016.11.012

Lawrence, B.A., Lishawa, S.C., Hurst, N., Castillo, B.T., Tuchman, N.C., 2017b. Wetland invasion by Typha × glauca increases soil methane emissions. Aquat Bot. 10.1016/j.aquabot.2016.11.012

Lawrence, B.A., Lishawa, S.C., Rodriguez, Y., Tuchman, N.C., 2016b. Herbicide management of invasive cattail (Typha × glauca) increases porewater nutrient concentrations. Wetl Ecol Manag 24. 10.1007/s11273-015-9471-x

Lishawa, S.C., Carson, B.D., Brandt, J.S., Tallant, J.M., Reo, N.J., Albert, D.A., Monks, A.M., Lautenbach, J.M., Clark, E., 2017. Mechanical harvesting effectively controls young typha spp. Invasion and unmanned aerial vehicle data enhances post-treatment monitoring. Front Plant Sci 8. 10.3389/fpls.2017.00619

Lishawa, S.C., Jankowski, K., Geddes, P., Larkin, D.J., Monks, A.M., Tuchman, N.C., 2014. Denitrification in a Laurentian Great Lakes coastal wetland invaded by hybrid cattail (Typha × glauca). Aquat Sci 76, 483–495. 10.1007/s00027-014-0348-5

Mitchell, M.E., Lishawa, S.C., Geddes, P., Larkin, D.J., Treering, D., Tuchman, N.C., 2011. Time-dependent impacts of cattail invasion in a great lakes coastal wetland complex, in: Wetlands. 10.1007/s13157-011-0225-0

Pieper, S., Dorken, M., Freeland, J., 2020a. Genetic structure in hybrids and progenitors provides insight into processes underlying an invasive cattail (Typha × glauca) hybrid zone. Heredity (Edinb) 124, 714–725. 10.1038/s41437-020-0307-y

Pieper, S., Dorken, M., Freeland, J., 2020b. Genetic structure in hybrids and progenitors provides insight into processes underlying an invasive cattail (Typha × glauca) hybrid zone. Heredity (Edinb) 124, 714–727. 10.1038/s41437-020-0307-y

Pieper, S.J., Freeland, J.R., Dorken, M.E., 2018. Coexistence of Typha latifolia, T. angustifolia (Typhaceae) and their invasive hybrid is not explained by niche partitioning across water depths. Aquat Bot 144, 46–53. 10.1016/j.aquabot.2017.11.001

Pieper, S.J., Nicholls, A.A., Freeland, J.R., Dorken, M.E., 2017. Asymmetric Hybridization in Cattails (Typha spp.) and Its Implications for the Evolutionary Maintenance of Native Typha latifolia. Journal of Heredity 108, 479–487. 10.1093/jhered/esx036

Shapro, A., 2022. Understanding morphological responses of Typha (cattail)species to nutrient pollution (MSc thesis). University of Northern Iowa, https://scholarworks.uni.edu/etd/1251.

Smith, S., 1967. Experimental and natural hybrids in North American Typha (Typhaceae). American Midland Naturalist 78, 257–287. 10.2307/2485231

Smith SG, 2020. Typhaceae [WWW Document]. Flora of North America.

Smith, S.G., 1967. Experimental and Natural Hybrids in North American Typha (Typhaceae). American Midland Naturalist 78, 257. 10.2307/2485231

Snow, A.A., Travis, S.E., Wildova, R., Fer, T., Sweeney, P.M., Marburger, J.E., Windels, S., Kubatova, B., Goldberg, D.E., Mutegi, E., 2010. Species-specific SSR alleles for studies of hybrid cattails (Typha latifolia x T. angustifolia; Typhaceae) in North America. Am J Bot 97, 2061–2067. 10.3732/ajb.1000187

Solberg, K.L., Higgins, K.F., 1993. Effects of Glyphosate Herbicide on Cattails, Invertebrates, and Waterfowl in South Dakota Wetlands. Wildl Soc Bull. 10.2307/3782869

Tangen, B.A., Bansal, S., Freeland, J.R., Travis, S.E., Wasko, J.D., McGonigle, T.P., Goldsborough, L.G., Gow, K., Marburger, J.E., Meier, J.A., 2022a. Distributions of native and invasive Typha (cattail) throughout the Prairie Pothole Region of North America. Wetl Ecol Manag 30, 1–17. 10.1007/s11273-021-09823-7

Tangen, B.A., Bansal, S., Freeland, J.R., Travis, S.E., Wasko, J.D., McGonigle, T.P., Goldsborough, L.G., Gow, K., Marburger, J.E., Meier, J.A., 2022b. Distributions of native and invasive Typha (cattail) throughout the Prairie Pothole Region of North America. Wetl Ecol Manag 30. 10.1007/s11273-021-09823-7

Tisshaw, K., Freeland, J., Dorken, M., 2020. Salinity, not genetic incompatibilities, limits the establishment of the invasive hybrid cattail Typha × glauca in coastal wetlands. Ecol Evol 10, 12091–12103. 10.1002/ece3.6831

Travis, S.E., Marburger, J.E., Windels, S., Kubátová, B., 2010a. Hybridization dynamics of invasive cattail (Typhaceae) stands in the Western Great Lakes Region of North America: A molecular analysis. Journal of Ecology 98, 7–16. 10.1111/j.1365-2745.2009.01596.x

Travis, S.E., Marburger, J.E., Windels, S., KubÃ¡tovÃ¡, B., 2010b. Hybridization dynamics of invasive cattail (Typhaceae) stands in the Western Great Lakes Region of North America: a molecular analysis. Journal of Ecology 98, 7–16. 10.1111/j.1365-2745.2009.01596.x

Tuchman, N.C., Larkin, D.J., Geddes, P., Wildova, R., Jankowski, K., Goldberg, D.E., 2009a. Patterns of environmental change associated withTypha xglauca invasion in a Great Lakes coastal wetland. Wetlands 29, 964–975. 10.1672/08-71.1

Tuchman, N.C., Larkin, D.J., Geddes, P., Wildova, R., Jankowski, K., Goldberg, D.E., 2009b. Patterns of environmental change associated withTypha xglauca invasion in a Great Lakes coastal wetland. Wetlands 29. 10.1672/08-71.1

Wasko, J.D., McGonigle, T.P., Goldsborough, L.G., Wrubleski, D.A., Badiou, P.H., Armstrong, L.M., 2022. Use of shoot dimensions and microscopic analysis of leaves to distinguish Typha latifolia, Typha angustifolia, and their invasive hybrid Typha xglauca. Wetl Ecol Manag 30. 10.1007/s11273-021-09836-2

Wendell, D.L., Huang, X., Gryspeerd, B., Freeland, J., 2021. A simple screen to detect hybrids between native and introduced Phragmites australis in the United States and Canada. J Great Lakes Res 47, 1453–1457. 10.1016/J.JGLR.2021.08.002

Ye, J., Coulouris, G., Zaretskaya, I., Cutcutache, I., Rozen, S., Madden, T.L., 2012. Primer-BLAST: a tool to design target-specific primers for polymerase chain reaction. BMC Bioinformatics 13. 10.1186/1471-2105-13-134

Zapfe, L., Freeland, J.R., 2015. Heterosis in invasive F-1 cattail hybrids (Typha x glauca). Aquat Bot 125, 44–47. 10.1016/j.aquabot.2015.05.004

